# Using semi-folded arenas to observe spontaneous emergence of collective trails in whole colonies of termites

**DOI:** 10.64898/2026.06.24.734206

**Authors:** Helder Hugo, Iain Douglas Couzin

**Author notes:** Corresponding author (HH).

## Abstract

Collective movement in social organisms emerges from local interactions and can generate large-scale spatial patterns of ecological relevance. In termites, trail formation is a well-known collective phenomenon, yet reproducing and recording its emergence under controlled laboratory conditions using whole colonies remains challenging. Existing laboratory approaches often rely on confined arenas or manually assembled subgroups, which can restrict movement and limit observation of colony-level dynamics. Here, we present a semi-folded arena designed for whole-colony observation of termite movement under controlled conditions. We developed a circular semi-folded arena that remained continuously connected to an intact nest and allowed individuals to move across a central observation surface while recirculating through a folded peripheral section. Using whole colonies of the Neotropical termite *Constrictotermes cyphergaster*, we recorded exploratory activity under baseline conditions, in the absence of added food or water. High-resolution video recordings were analysed using automated movement extraction to recover trajectories and visualise collective trail structure. Within the first 6 min of activity, collective trail structure was observed in 15 of the 16 colonies analysed. Under these conditions, the semi-folded setup captured early collective trail structure, visible as convergence of cumulative trajectories along shared routes radiating from the arena entrance region. Automated movement extraction was compatible with dense whole-colony recordings and yielded large quantities of positional data during the initial observation interval. Descriptive trajectory-based outputs, including speed distributions for workers and soldiers, showed that the recordings were suitable for recovery of fine-scale movement information. Repeatedly used routes were also often marked by visible dark traces on the paper lining by the end of the observations, providing a qualitative record of cumulative route use. The semi-folded arena provides a practical method for recording whole-colony termite movement under laboratory conditions while maintaining continuous nest access and avoiding manual transfer of individuals during trials. Rather than replacing conventional arena designs, this approach offers an additional methodological option for studying emergent movement patterns in species for which whole-colony observation is feasible. More broadly, it expands the experimental toolkit available for investigating colony-scale spatial organisation under controlled conditions.

## BACKGROUND

Understanding how collective movement emerges in social organisms depends on methods that allow behaviour to be observed under controlled conditions without obscuring the spatial structure of interactions. This is particularly challenging in systems where individuals operate within enclosed nests, under low light, or in dense aggregations that are difficult to record without disturbing the colony. From a movement ecology perspective, such constraints limit the study of how local interactions scale up into colony-level movement patterns and spatial organisation [1-4].

In eusocial insects, these challenges are especially evident. Colony members may remain inactive for extended periods, perform cooperative tasks within concealed nest spaces, or interact in large numbers under conditions that are difficult to standardise experimentally [2,5,6]. In termites, many ecologically relevant behaviours occur either within nests or along exposed or transient foraging routes, making them difficult to capture in ways that combine direct observation with experimental control [5,7]. As a result, methodological choices can strongly influence which aspects of collective movement become accessible to study.

Many laboratory studies with termites rely on standardised indoor arenas in which individuals or subsets of colonies are confined within enclosed spaces. Containers such as Petri dishes have been widely used to investigate a range of topics, including exploratory behaviour [8], aggression [9-11], sexually dimorphic movement [12], tandem running [13], infection-related responses [14], alarm behaviour [15], and caste differentiation [16]. These approaches have been highly informative, but they have generally focused on individual-level or small-group processes. Such designs are often well suited to eliciting specific interactions, yet their spatial scale can constrain the expression of distributed, colony-level dynamics that are central to the study of self-organisation and collective behaviour [1-3].

At the same time, termite arena design is often adapted to the biological context and to the specific question being addressed. In some studies, enclosed containers such as Petri dishes have been used deliberately to increase encounter rates under particular ecological conditions, such as interspecific interactions [17] or mating behaviour [18]. In others, more specialised designs have been developed to accommodate the biology of different species and to address particular behavioural questions. For instance, interconnected planar arenas have been used to study subterranean termites [19], glass tubes have enabled direct observation of behavioural responses to air movement [20], and custom arenas have been used in tunnelling [21], construction [22], or vibroacoustic assays [23]. Together, these examples show that experimental design in termites is closely tied to biological context and research aim, and that methods optimised for local interactions do not necessarily lend themselves to observing whole-colony movement patterns at broader spatial scales [2-4].

Whole-colony observations are particularly valuable in the study of self-organisation and collective behaviour because they allow local interactions to be examined within the broader social and spatial context in which they occur. Such methods have been implemented successfully in other eusocial taxa, especially within Hymenoptera. In bees, for example, whole-colony observation systems have been used to investigate spatial organisation within the nest [24], social interactions and network structure [25], and emergent colony-level dynamics [26]. Likewise, in ants, colony-scale experimental designs have been used to study spatial fidelity and colony organisation [27], collective exploration and trail pattern formation [28,29], persistent trail use [30], adaptive transport networks under changing resource conditions [31], exploration of novel environments [32], and wall-following in a social context [33].

In termites, however, comparable whole-colony studies remain less common. Recent movement studies at this scale have focused mostly on invasive subterranean species and applied questions related to foraging or bait interception [19,34]. Studies addressing broader behavioural questions also exist, including analyses of trail dynamics [35], but remain relatively uncommon. This relative scarcity reflects both the practical difficulty of working with whole colonies in many termite species and the limited use of laboratory setups suited to recording large numbers of termites moving simultaneously under controlled conditions. For species in which colony-scale observation is feasible, there remains a need for methods that allow colony-level movement patterns to be documented while maintaining direct nest access.

Here, we present a semi-folded arena for whole-colony observation of termite movement under laboratory conditions. The system remains continuously connected to intact nests containing whole colonies and provides a central observation surface from which individuals can recirculate through a folded peripheral section. Using the Neotropical termite *Constrictotermes cyphergaster* as a model, we evaluate the method as a proof of concept by specifically asking whether the setup (1) supports the emergence of observable colony-level trail structure under baseline conditions and (2) produces recordings compatible with extraction of movement data from dense whole-colony activity. This study aims to introduce this approach as an additional methodological option for studies in which whole-colony observation is both feasible and desirable, rather than to benchmark it against all alternative designs.

## MATERIALS AND METHODS

### Biological species, study site and sampling

*Constrictotermes cyphergaster* (Termitidae: Nasutitermitinae) is a Neotropical termite widely distributed across South America and particularly common in the drylands of Brazil [36]. It forages on exposed surfaces without the protection of covered galleries, and mature colonies typically occupy arboreal nests attached to tree trunks [37-38] (Figure 1A-B). The developmental pathway of the apterous line has been described in detail, mentioning the worker caste as monomorphic [39], although recent published accounts differ in describing them dimorphic [40]. Unlike workers, soldiers are differentiated through a presoldier stage [39] and bear a characteristic nasus-based chemical defence (Figure 1C) rather than functional biting mandibles [6,41]. Despite these differences, individuals of both castes in the colonies observed here were of broadly similar size (body lengths ranging from 4 to 5 mm), with similar detected image footprints in video frames (99.2 ± 28.8 pixels per individual). In addition to its conspicuous external foraging behaviour, *C. cyphergaster* has attracted increasing attention because its nests frequently harbour associated termitophiles, including inquiline termites [17], rove beetles [42,43], and potter wasps [44]. These features make the species a suitable model for developing and testing methods for colony-level observation under laboratory conditions, including whole-colony assays where intact colonies are available.

**Figure 1.**
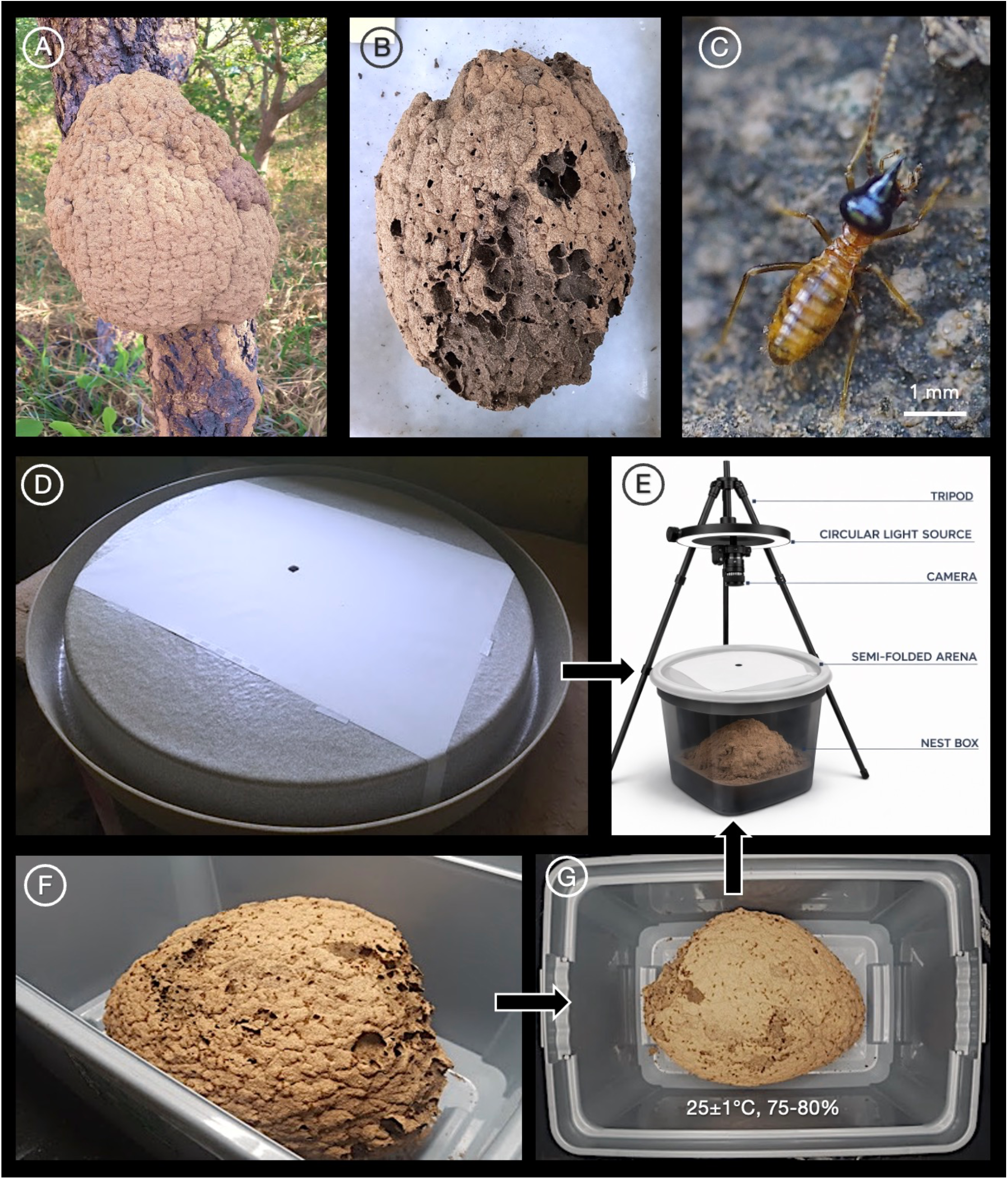
Study system and experimental setup. (A,B) Arboreal nests of *C. cyphergaster*. (C) Soldier of *C. cyphergaster*; visual scale = 1mm (D) Semi-folded circular arena lined with DIN A3 tracing paper; internal diameter of the central observation surface = 490 mm, external diameter including the folded section = 560 mm, groove depth = 45 mm. (E) Schematic view of the setup; the nest-box wall is shown as transparent to aid visualisation. (F,G) Nest box used to maintain whole colonies under controlled temperature and humidity (25 ± 1 °C, 75-80% relative humidity).

We collected 16 nests of *C. cyphergaster* in the Brazilian Cerrado near the municipality of Divinópolis, Minas Gerais, south-eastern Brazil (20°08′20″S, 44°53′02″W), in an area characterised by tropical savanna vegetation with dry winters [45,46]. Sampling was conducted at the beginning of the wet season, in November 2019 and November 2020. Entire nests were detached manually from tree trunks by a single researcher (HH). *C. cyphergaster* nests are typically attached to the substrate by a relatively narrow basal section, which allows detachment with limited structural damage [36-39] (Figure 1A-B).

Immediately after collection, each nest was placed in an individual plastic nest box (**Figure 1F**) and transported to the experimental room. All colonies were maintained in their original nest structures throughout the study. Inside the nest box, the exposed side of the nest, corresponding to the original attachment point to the tree, was positioned facing downward against the box floor (**Figure 1E**). This substantially closed galleries opened during sampling, although minor gaps could remain because of nest irregularities.

### Semi-folded geometry and peripheral circulation pathway

To provide a large observation surface while avoiding abrupt vertical walls in the main movement area, we developed a circular arena with semi-folded geometry (Figure 1D). Our arenas were manufactured from plastic using press-forming and comprised a flat central platform surrounded by a rounded peripheral fold. The internal diameter of the observable central surface was 490 mm, and the external diameter, including the folded section, was 560 mm. The peripheral fold formed a continuous U-shaped groove approximately 45 mm deep around the arena rather than a sharp corner or wall. The outermost wall of the folded section was smooth enough to prevent *C. cyphergaster* termites from escaping under our experimental conditions. By contrast, the remaining arena surfaces were left slightly textured to allow termites to move freely across the arena. The transition between the central platform and the peripheral fold was intentionally rounded to avoid a sharp edge that could influence termite movement or hinder transitions between the observable surface and the U-shaped groove.

Arenas remained continuously connected to a colony nest box throughout each trial (Figure 1E). Termites reached the arena from below through an opening aligned with the centre of the platform. A flexible plastic bridge (20 cm long, 2 cm wide) linked the exposed lower surface of the nest to the entrance beneath the arena, allowing individuals to emerge onto the observation surface and later recirculate through the peripheral fold without manual transfer during observations. Individuals leaving the central area and descending into the folded section could subsequently climb the U-shaped groove and re-enter the observation area, allowing colonies to circulate continuously through the system. The folded geometry is therefore a defining feature of the design, because individuals leaving the central observation surface can continue moving through the groove and later re-enter the observable area rather than encountering an abrupt lateral wall at the edge of the main surface.

Before each whole-colony observation, the arena was lined with tracing paper (Figure 1D). In preliminary tests, we evaluated several candidate surface materials, including acrylic, frosted glass, and different paper types. Tracing paper provided the most reliable locomotor performance under our conditions, whereas lighter papers warped under ambient moisture and smoother rigid surfaces were less practical at the dimensions required for whole-colony observations. We therefore used DIN A3 tracing paper (297 × 420 mm) with a grammage of at least 190 g/m^2^, which remained stable during the recording period. The arena dimensions were chosen so that DIN A3 tracing paper could be fitted securely to the central region while preserving access to the peripheral fold. The paper served two purposes. First, it provided a consistent walking surface that supported stable locomotion and image contrast during tracking. Second, it retained visible traces after repeated route use, allowing qualitative inspection of cumulative route marking at the end of each observation. Because termites sometimes followed the exposed edge of the paper, the narrow peripheral zone adjacent to the paper margin was excluded from the main trackable area, as shown in Figure 2.

**Figure 2.**
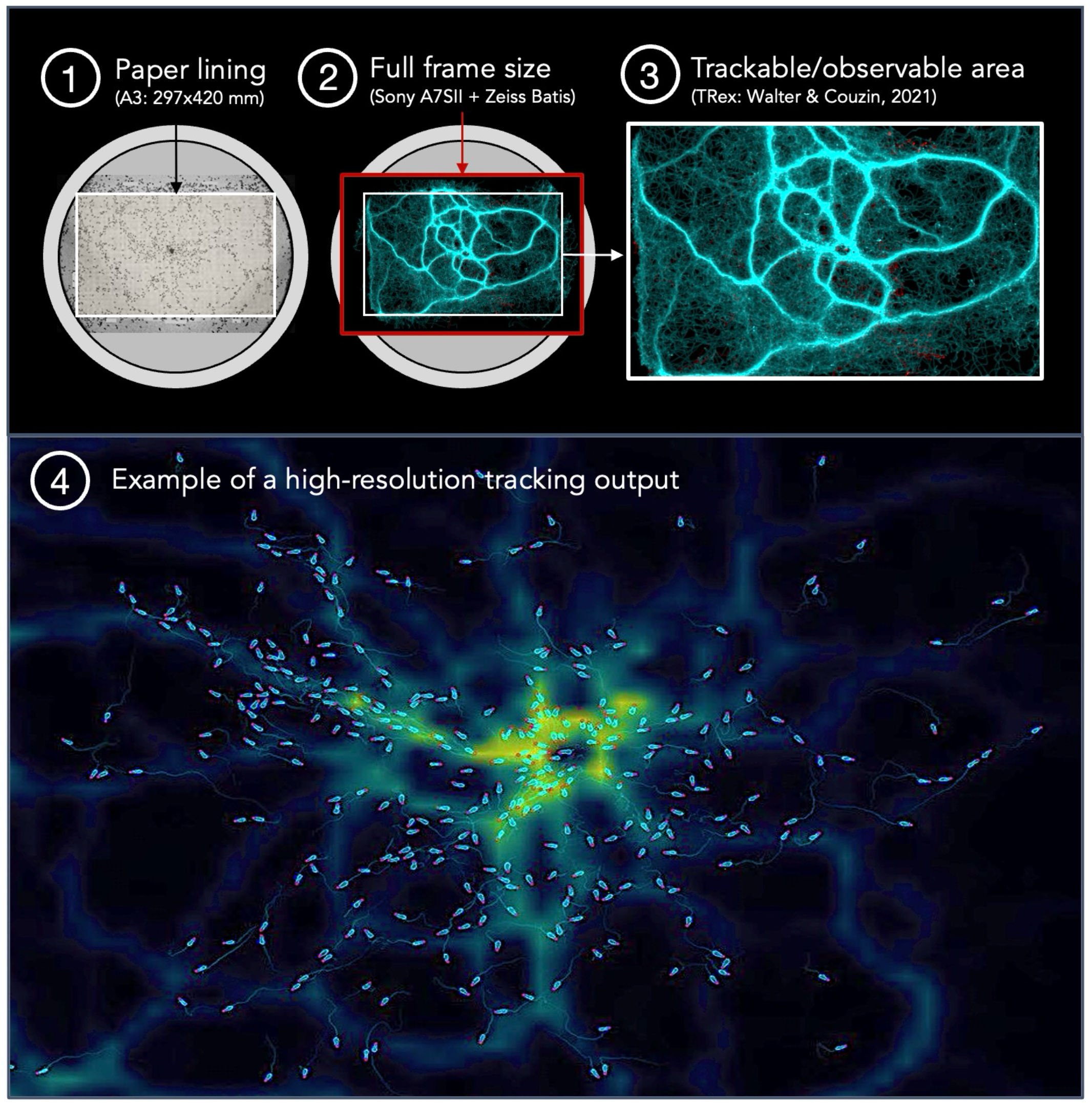
Movement data acquisition from high-resolution recordings. (1-3) Schematic overview of the paper-lined semi-folded arena, the full camera frame, and the observable (trackable) area defined for automated trajectory extraction. (4) Example of a high-resolution tracking output, combining individual detections with a spatial visualisation of movement density across the observation area.

### Recording setup and environmental conditions

Each recording used a unique termite nest containing an intact colony. Whole-colony activity was video recorded for 2 h 30 min, but the present study focused on the first 6 min after termites began to emerge onto the arena. This interval corresponded to 9,000 frames per colony and was used because it captured the initial phase of arena use while allowing consistent comparison across colonies.

Observations were recorded under visible light in 4K resolution at 25 frames per second using a Sony Alpha 7S II camera fitted with a Zeiss Batis 25 mm f/2 lens. Illumination was provided by a ring light (NEEWER RL-18, 18-inch LED, 55 W, 240 LEDs, bi-colour output adjustable from 3200 K to 5600 K, CRI 95, 0-100% dimmable; no discrete peak wavelength reported by the manufacturer) positioned concentrically around the lens to provide diffuse and relatively uniform lighting across the arena surface. The camera was mounted vertically 50 cm above the arena (Figure 1E). Focus was adjusted manually using the camera zoom-assist function to maximise sharpness on the central observation plane. Before each recording, the camera axis and field of view were aligned using calibration marks and a printed graphical scale attached to the paper lining. This ensured that the full observation surface was visible with a small margin and allowed conversion from image coordinates to real-world distances during subsequent movement extraction.

Because termites are highly sensitive to desiccation, reduced ambient moisture can rapidly impair their activity and survival, particularly in non-drywood species [47-50]. Stable temperature and humidity were therefore maintained between nest collection and arena observation. Colonies were kept inside their original nests until observations began, all experimental work was completed within 24 h of field collection, and relative humidity inside the nest boxes was maintained at 75-80%, with nest box and room temperature held at 24-26 °C (Figure 1F-G). These conditions were sufficient to preserve activity throughout the recording period.

### Proof-of-concept validation and movement data extraction

Videos were analysed using automated tracking methods suitable for static backgrounds and standardised lighting [51]. Movement data were extracted with TRex, an open-source software for multi-animal tracking from video recordings [52]. The analysis covered the first 6 min of each recording, corresponding to 9,000 frames per colony and 144,000 frames across the 16 colonies included in the dataset.

The methodological proof of concept evaluated two criteria. First, whether the semi-folded arena supported observable collective trail structure during the initial observation interval. Second, whether the recordings yielded movement data that could be extracted under dense whole-colony activity. For the first criterion, collective trail structure was scored as present when cumulative trajectories showed repeated convergence of multiple individuals along shared routes radiating from the arena entrance region within the first 6 min. For the second criterion, extraction output was summarised for each colony using total detections, mean detections per frame, and maximum detections per frame.

Extracted movement data were used to generate whole-colony movement visualisations (Figure 3A-O), caste-specific speed distributions (Figure 4A-B), and representative individual trajectories as qualitative examples of the image-based outputs obtainable from the recordings (Figure 4C-D). Because dense local occlusion sometimes interrupted continuous identity assignment in heavily occupied regions, validation was based on movement detections and reconstructed trajectory patterns rather than on validated long-term identity persistence for every visible individual. Behavioural modelling based on these data is possible but falls outside the scope of the present methodological study.

**Figure 3.**
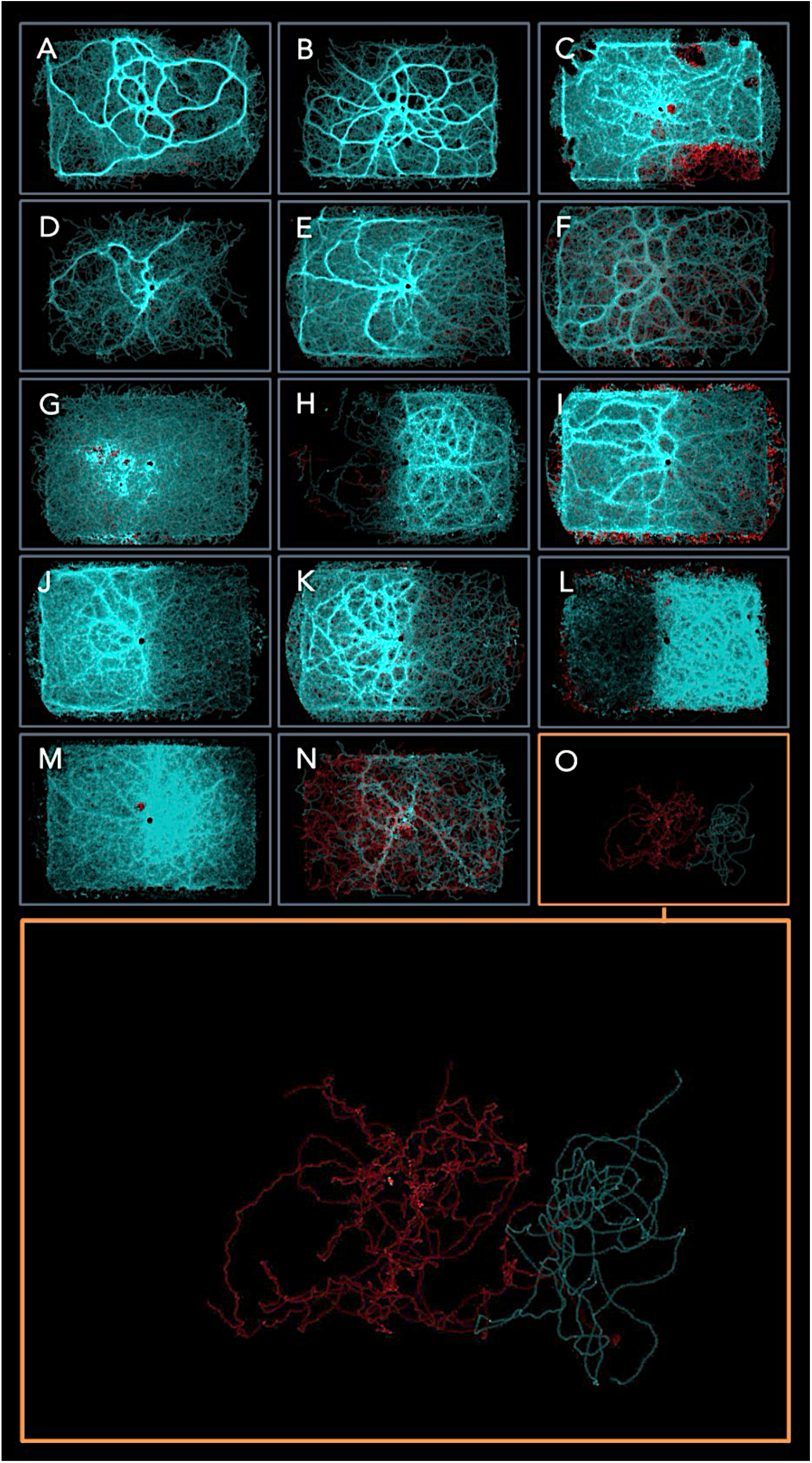
Spontaneous emergence of collective trails. **(A-O)** Cumulative trajectories from whole-colony observations in semi-folded arenas. Each panel shows trajectories extracted from the first 9,000 frames (6 min) of a different termite colony, with worker trajectories in blue and soldier trajectories in red. The panels illustrate colony-level route structure during the initial 6 min of observation. Because the pattern in colony O occupied a smaller area, it is also shown enlarged for clarity (bottom).

**Figure 4.**
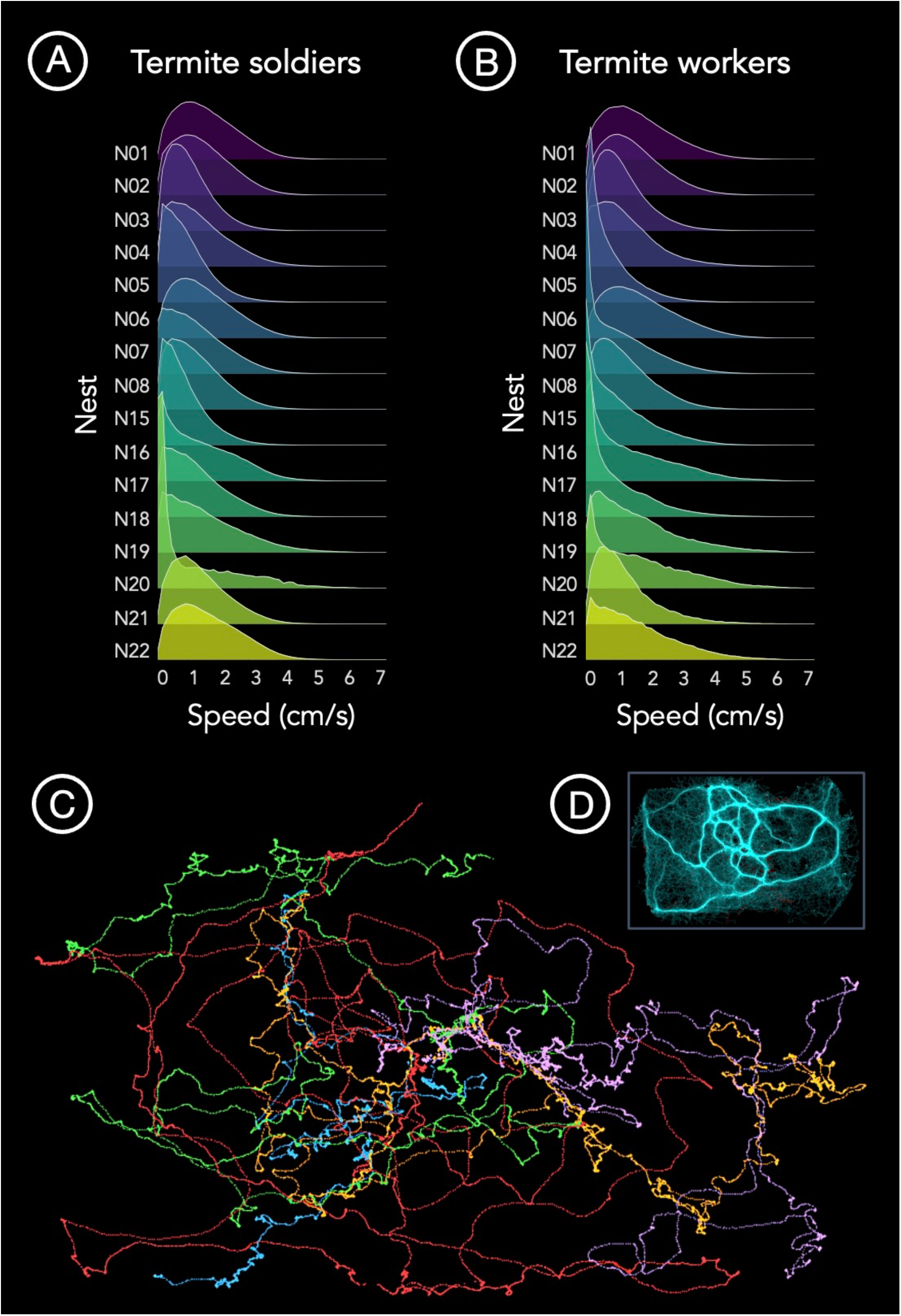
Descriptive speed profiles and examples of trajectory-based outputs. (A,B) Kernel density estimates of speed for termite soldiers (A) and workers (B) from each whole-colony observation, based on the first 6 min of movement extraction (9,000 frames). Nests collected in 2019 are labelled N15-N22, and those collected in 2020 are labelled N01-N08. (C) Example trajectories of individual termites, each shown in a different colour. (D) Cumulative trajectory pattern for the whole-colony recording used to illustrate the individual trajectories shown in panel C.

## RESULTS

### Collective trail structure under baseline conditions

Collective trail structure, as defined in Materials and Methods, was observed within the first 6 min in 15 of the 16 colonies analysed (Table 1). In these colonies, cumulative trajectories showed repeated convergence of multiple individuals along shared routes radiating from the arena entrance region during the initial observation interval. One colony did not meet this criterion within the first 6 min and was therefore not included in the trajectory panel used to illustrate early trail structure. Detection-based movement output was recovered from all colonies. Across recordings, total detections ranged from 153,383 to 12,853,513 per colony, mean detections per frame ranged from 17.0 to 1,428.2, and maximum detections per frame ranged from 22 to 2,339 (Table 1). These values represent detection events rather than counts of unique individuals and show that the recording workflow yielded large quantities of movement data during dense whole-colony activity.

**Table 1.**
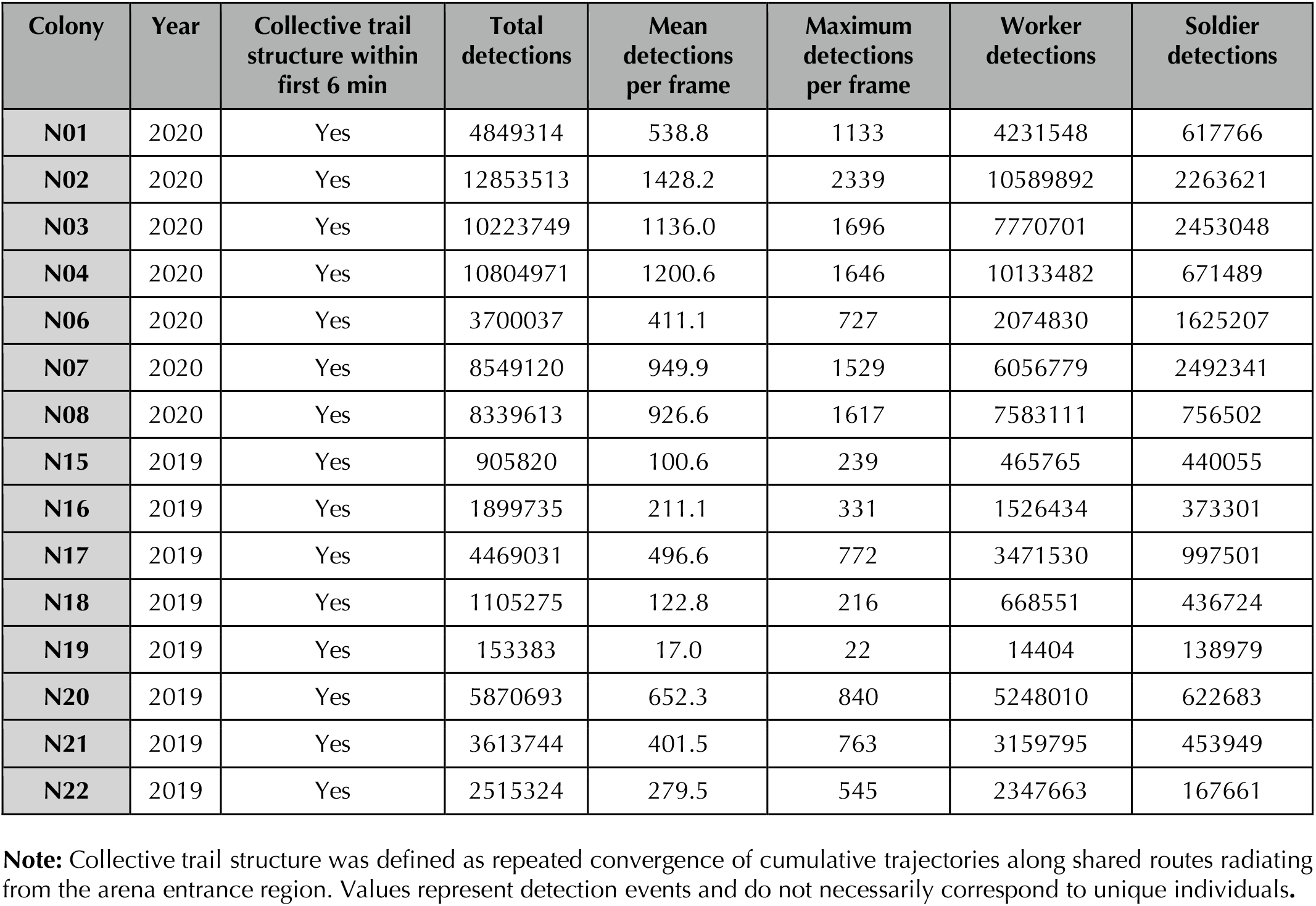
Proof-of-concept validation metrics. Measurements were derived from the initial observation period (first 6 min) of whole-colony recordings.

| Colony | Year | Collective trail structure within first 6 min | Total detections | Mean detections per frame | Maximum detections per frame | Worker detections | Soldier detections |
| --- | --- | --- | --- | --- | --- | --- | --- |
| N01 | 2020 | Yes | 4849314 | 538.8 | 1133 | 4231548 | 617766 |
| N02 | 2020 | Yes | 12853513 | 1428.2 | 2339 | 10589892 | 2263621 |
| N03 | 2020 | Yes | 10223749 | 1136.0 | 1696 | 7770701 | 2453048 |
| N04 | 2020 | Yes | 10804971 | 1200.6 | 1646 | 10133482 | 671489 |
| N06 | 2020 | Yes | 3700037 | 411.1 | 727 | 2074830 | 1625207 |
| N07 | 2020 | Yes | 8549120 | 949.9 | 1529 | 6056779 | 2492341 |
| N08 | 2020 | Yes | 8339613 | 926.6 | 1617 | 7583111 | 756502 |
| N15 | 2019 | Yes | 905820 | 100.6 | 239 | 465765 | 440055 |
| N16 | 2019 | Yes | 1899735 | 211.1 | 331 | 1526434 | 373301 |
| N17 | 2019 | Yes | 4469031 | 496.6 | 772 | 3471530 | 997501 |
| N18 | 2019 | Yes | 1105275 | 122.8 | 216 | 668551 | 436724 |
| N19 | 2019 | Yes | 153383 | 17.0 | 22 | 14404 | 138979 |
| N20 | 2019 | Yes | 5870693 | 652.3 | 840 | 5248010 | 622683 |
| N21 | 2019 | Yes | 3613744 | 401.5 | 763 | 3159795 | 453949 |
| N22 | 2019 | Yes | 2515324 | 279.5 | 545 | 2347663 | 167661 |

**Note:** Collective trail structure was defined as repeated convergence of cumulative trajectories along shared routes radiating from the arena entrance region. Values represent detection events and do not necessarily correspond to unique individuals.

Cumulative trajectories from the 15 colonies that met the collective trail structure criterion within the first 6 min are shown in Figure 3. In these recordings, repeated use of shared routes produced visible colony-level path structure across the central observation surface. Because no food or water was added to the arena, these visualisations indicate that the semi-folded setup was sufficient to capture early route convergence under standardised laboratory conditions.

### Movement data from dense whole-colony activity

In addition to whole-colony movement visualisations, extracted movement data were used to generate caste-specific speed distributions (Figure 4A-B), as well as representative individual trajectories (Figure 4C-D). These outputs are included as examples of the type of movement information recoverable from the recordings rather than as the basis for formal behavioural inference. In both castes, speed distributions were typically unimodal, with many observations concentrated at relatively low speeds and a right-skewed tail extending towards higher values. Considerable within-caste variation was evident across colonies, and the distributions of workers and soldiers overlapped broadly across the dataset. Together, these descriptive outputs show that the protocol can recover fine-scale movement information from dense whole-colony recordings.

At the end of whole-colony observations, repeatedly used routes were often marked by visible dark traces on the paper lining (Figure 5). These markings were spatially consistent with routes that had been used repeatedly during the recording period and provided a qualitative record of cumulative route use on the arena surface. The chemical identity and behavioural function of these deposits were not examined here.

**Figure 5.**
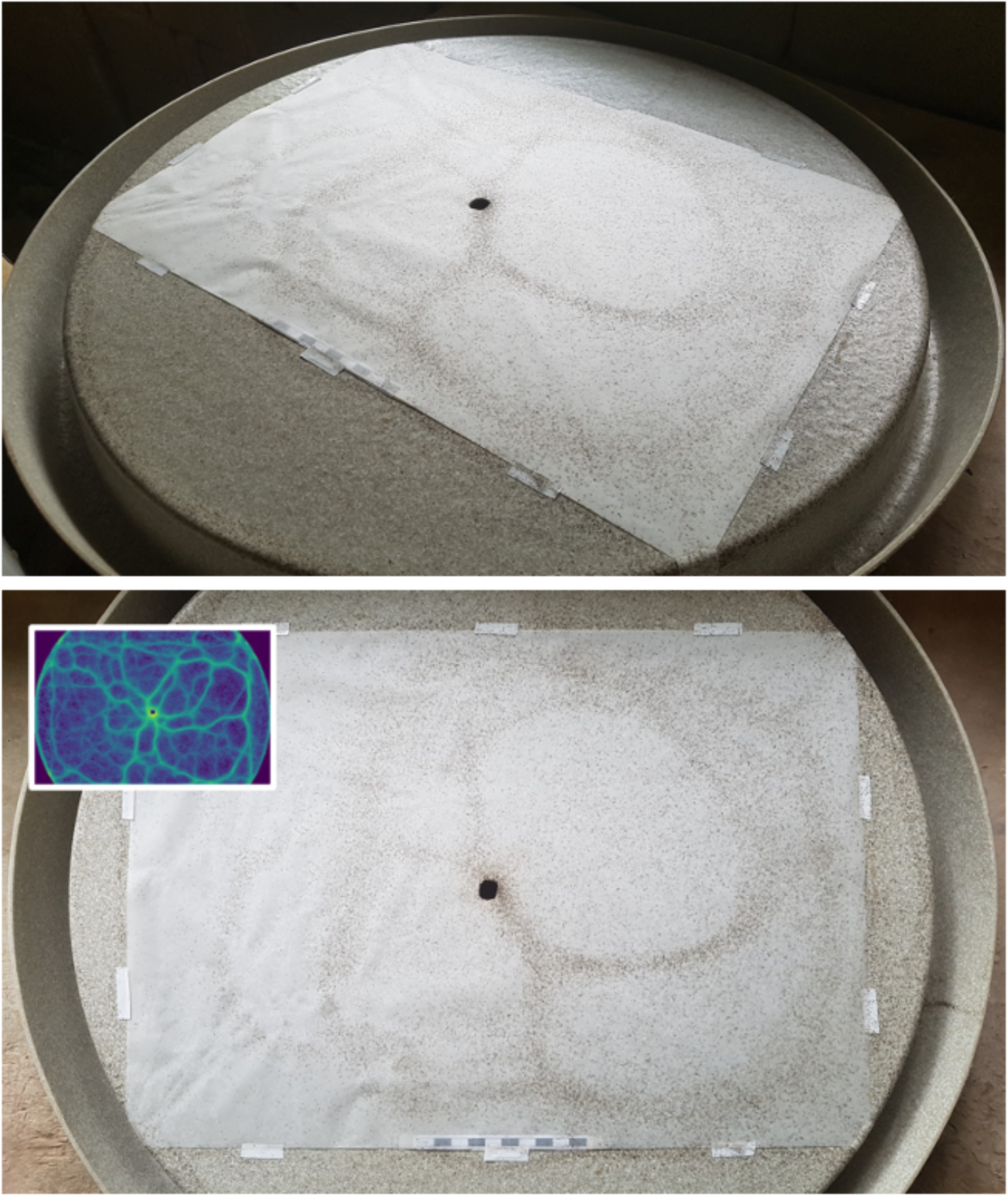
Visible traces along repeatedly used routes. Top and bottom panels show the semi-folded arena after whole-colony observations, with darker traces along frequently used paths on the paper lining. These traces are consistent with repeated route use and retention of deposited material, previously associated with faecal marking in *C. cyphergaster* [54]. Inset: corresponding spatial density map of movement.

## DISCUSSION

We introduce a semi-folded arena and recording workflow for observing colony-scale termite movement under controlled laboratory conditions. In *C. cyphergaster*, the setup supported early collective trail structure within the first 6 min in 15 of 16 colonies and produced dense movement data that could be extracted automatically from whole-colony recordings. The contribution of the present study is therefore methodological. It shows that this configuration can be used to document early route convergence and generate trajectory-based outputs from colony-scale activity while maintaining continuous connection between the observation surface and the original nest.

### Advantages of the method

A central feature of the method is the semi-folded geometry. Unlike a conventional flat arena bounded by abrupt lateral walls, the folded peripheral section provides a continuous pathway through which termites leaving the main observation surface can continue moving and later re-enter the observable area. Under the conditions tested here, this geometry supported circulation through the apparatus while preserving a large central surface suitable for imaging and movement extraction. The main value of the design lies in this combination of recirculation and observability, rather than in whole-colony observation alone.

Continuous connection to the intact nest box provided a second practical advantage. Colonies could enter and leave the arena without manual transfer during the recording period, reducing disturbance at the onset of activity and preserving the immediate social context from which movement on the arena emerged. This feature may be useful in studies where colony-scale spatial organisation is sensitive to local density, recruitment, or nest access under controlled conditions.

The value of the setup also depended on the recording workflow. Standardised lighting, fixed camera geometry, and automated movement extraction allowed dense colony activity to be converted into cumulative trajectories, representative path reconstructions, and descriptive movement distributions. The method should therefore be viewed as the combination of arena design and imaging workflow, rather than as a purely structural modification of the arena.

The paper-lined surface offered an additional practical benefit by preserving visible route traces over the course of an extended observation. In *C. cyphergaster*, persistent route marking has previously been associated with faecal deposits [54], and trail communication in Nasutitermitinae termites more broadly may also involve sternal gland pheromones [53-56]. We did not examine the chemical identity or behavioural function of the deposits observed here, but the retained traces show that the paper-lined surface can preserve a cumulative spatial record of repeated route use.

### Limitations of the method

Like most experimental designs, the method also has its limitations. First, the arena remains an artificial two-dimensional environment and does not reproduce the structural complexity of natural termite habitats, including bark surfaces, canopy routes, tunnels, or three-dimensional nesting substrates. It should therefore be regarded as a controlled observational platform rather than as a direct representation of field conditions.

Second, the semi-folded arena was not compared directly with conventional closed arenas or other laboratory designs. The present study therefore does not permit formal inference about relative performance across arena types. The results show only that the semi-folded configuration provided workable conditions for recording early route convergence and extracting dense movement data under the parameters used here.

A further limitation concerns generality across taxa. The protocol was developed for *C. cyphergaster*, an arboreal species that forages on exposed surfaces [7,36-38]. Species that are strongly photophobic, highly cryptic, or adapted to confined subterranean movement may require different surfaces, lighting regimes, or experimental layouts. In addition, dense local occlusion can interrupt identity continuity in heavily occupied areas, so the present proof-of-concept validation rests on detection-based outputs and reconstructed trajectory patterns rather than on validated long-term identity persistence for every visible individual. Over longer observation periods, repeated circulation through the folded section may also promote peripheral route reuse, local accumulation, or recurrent looping within parts of the groove. These longer-term dynamics were outside the scope of the present study, which focused on the initial 6 min of arena use.

### Scope, future applications, and conclusion

Taken together, our results indicate that semi-folded arenas can support whole-colony observation of early shared route structure under controlled laboratory conditions while remaining compatible with automated movement extraction. In the species and experimental context examined here, *C. cyphergaster*, the setup allowed colony-level route structure to be documented during the first minutes of activity and yielded descriptive trajectory-based outputs from dense whole-colony observations.

At the same time, the scope of the present paper is intentionally limited. The study does not benchmark the semi-folded arena against conventional closed arenas or other laboratory designs. Also, it not an attempt to provide a full behavioural analysis of trail development, caste function, or route reinforcement, something that remains to be further explored. The main value of the method lies not in demonstrating superiority over alternative approaches, but in showing that this configuration provides workable conditions for recording colony-scale movement in a controlled setting while maintaining continuous connection to the original nest.

Within those limits, an approach using semi-folded arenas may be useful in future studies seeking to link whole-colony spatial organisation with trajectory-level movement data under controlled conditions. This may include work on route formation, caste-specific spatial structure, or the temporal development of collective patterns, provided that such questions are addressed with designs and analyses tailored to those aims. More broadly, our method expands the set of experimental options available for studying collective movement in termite species for which whole-colony observation is feasible and biologically meaningful.

## ETHICS STATEMENT

This study was conducted under the internal ethical oversight of the Max Planck Institute of Animal Behavior (MPIAB) and in line with the Best Practice Guidelines for Research of the Max Planck Society. Fieldwork and observations were carried out in Brazil under the relevant Brazilian scientific collection authorisations issued through SISBIO/ICMBio, in accordance with Instrução Normativa ICMBio nº 03/2014, and with permission from landowners to access sampling sites near the municipality of Divinópolis, Minas Gerais, Brazil. The research did not involve endangered or protected species, and no genetic resources were accessed.

## ACKNOWLEDGEMENTS

We thank Prof. Angelo Fonseca, Katia C. D. Santos, and Julio H. Santos for their valuable support during field expeditions. We are also grateful to Dr Tristan Walter, lead developer of TRex, for assistance with video processing and tracking, and for making the software openly available. We thank the Department of Entomology at the Universidade Federal de Viçosa and members of the Laboratory of Termitology for logistical support in Brazil throughout different stages of the study. We acknowledge the Department of Zoology at the University of Cambridge for providing a supportive research environment and access to facilities during the final development of this study. Finally, we thank the organisers and participants of the workshop *Mathematics of Movement*, held at the Isaac Newton Institute for Mathematical Sciences in Cambridge, for constructive discussions that helped shape this work.

## FUNDING

HH received support from the Deutscher Akademischer Austauschdienst (DAAD-57214224), the International Max Planck Research School for Quantitative Behaviour, Ecology and Evolution (formerly the International Max Planck Research School for Organismal Biology), the Max Planck Institute of Animal Behavior, and the Max Planck Society. IDC received support from the National Science Foundation (IOS-1355061, DBI-2021795), the Office of Naval Research (N00014-19-1-2556), Horizon 2020 (860949, 101098722), the Deutsche Forschungsgemeinschaft under Germany’s Excellence Strategy (EXC 2117-422037984), and the Max Planck Society.

## DATA ACCESSIBILITY STATEMENT

Representative visual outputs and methodological details supporting the findings of this study are provided within the article. Non-digital material associated with species identification, including voucher specimens, is deposited in the Isoptera collection of the Universidade Federal de Viçosa, Minas Gerais, Brazil (UFV). Processed data underlying the methodological validation presented here may be made available by the corresponding author upon reasonable request.

## Notes

### Competing Interest Statement

The authors have declared no competing interest.

